# Localized activation of ependymal progenitors induces EMT-mediated glial bridging after spinal cord injury

**DOI:** 10.1101/2020.02.18.955401

**Authors:** Lili Zhou, Brooke Burris, Ryan Mcadow, Mayssa H. Mokalled

**Affiliations:** Department of Developmental Biology, Washington University School of Medicine

**Keywords:** Spinal cord injury, glial bridging, zebrafish, regeneration, Yap, Ctgf, EMT

## Abstract

Unlike mammals, adult zebrafish undergo spontaneous recovery after major spinal cord injury. Whereas scarring presents a roadblock for mammalian spinal cord repair, glial cells in zebrafish form a bridge across severed spinal cord tissue to facilitate regeneration. Here, we performed FACS sorting and genome-wide profiling to determine the transcriptional identity of purified bridging glia. We found that Yap-Ctgf signaling activates epithelial to mesenchymal transition (EMT) in localized niches of ependymal cells to promote glial bridging and regeneration. Preferentially activated in early bridging glia, Yap is required for the expression of the glial bridging factor Ctgfa and for functional spinal cord repair. Ctgfa regulation is controlled by an injury responsive enhancer element that drives expression in early bridging glia after injury. Yap-Ctgf signaling activates a mesenchymal transcriptional program that drives glial bridging. This study revealed the molecular signatures of bridging glia and identified an injury responsive gene regulatory network that promotes spinal cord regeneration in zebrafish.

## INTRODUCTION

Traumatic spinal cord injuries (SCI) are incurable conditions that require long-term therapeutic, rehabilitative, and psychological interventions. Following SCI, primary neuronal loss triggers cascades of secondary damages in the mammalian spinal cord (SC). Scarring, neurotoxicity, and inherent resistance to axon growth and neurogenesis continue to present major hurdles to SC regenerative medicine.

In contrast to mammals, elevated regenerative capacity enables adult zebrafish to reverse paralysis within 6-8 weeks of complete SC transection [1, 2]. Zebrafish elicit efficient pro-regenerative injury responses including axon regrowth, neurogenesis, and absence of scarring [2–4]. These regenerative processes distinguish the zebrafish SC from mammalian SCs and enable its natural repair post-injury. Following SC transection in zebrafish, a group of specialized glial cells bridge the SC, and axons are believed to regrow along these bridging glia [1, 2]. Glial bridging is a natural regenerative mechanism that could be deployed to support axon growth across species. Translating this regenerative process into regenerative therapies requires a comprehensive understanding of the cell fates and molecular mechanisms that lead to the emergence of bridging glial cells after SCI.

We previously identified *connective tissue growth factor a* (*ctgfa*) as a pro-regenerative bridging factor in zebrafish [1]. Ctgf is a secreted, extracellular molecule that elicits various cellular responses including proliferation and differentiation. The spatiotemporal pattern of *ctgfa* expression correlates with glial bridge formation in zebrafish. Following SCI, *ctgfa* expression is first broadly induced within ependymo-radial glial cells (ERGs), which line the central canal proximal to both ends of the lesion. During subsequent steps of regeneration, *ctgfa* transcripts localize to bridging glial cells and surrounding cells at the lesion core, and to ventral ERGs that are thought to give rise to bridging glia. Genetic mutants in zebrafish *ctgfa* demonstrated its requirement during bridging and regeneration; whereas genetic and pharmacologic *ctgfa* overexpression proved sufficient to promote bridging and functional SC repair. These findings implicated Ctgf as a central player in SC bridging, and importantly, enabled us to generate a battery of Ctgf-based tools to decipher cell identities and molecular mechanisms during glial bridging in zebrafish.

The transcriptional co-activators Yes-associated protein (Yap) and transcriptional co-activator with PDZ-binding motif (Taz) are downstream co-activators of Hippo signaling, regulating cell plasticity and organ growth during development and regeneration. Yap and Taz lack DNA-binding domains and control transcription by association with the Tead DNA-binding transcription factors to promote cell proliferation and maintenance of stem cell fate [5, 6]. Ctgf is a direct target of Yap/Taz signaling in multiple tissue contexts, and it remains to be determined whether Yap signaling is activated and whether Yap activation regulates Ctgf expression and glial bridging after SCI in zebrafish.

Unlike CNS injuries, peripheral nerve injuries elicit pro-regenerative responses in mammals. Axon regrowth and myelination of regenerating axons enable functional repair after nerve injury. Glial cell injury responses are central to this regeneration process and involve the activation of a repair program by differentiated Schwann cells. Following transection injuries that generate a gap in the mammalian nerve, repair Schwann cells migrate into the lesion site, bridge the severed ends of the nerve, and guide regenerating axons through the nerve bridge to support distal innervation [7, 8]. Repair Schwann cells execute these regenerative functions by upregulating EMT-associated genes during nerve regeneration, which result in partial cellular reprogramming from an epithelial cell fate to a more plastic mesenchymal fate [9, 10]. The Twist, Zeb, Snail, and Slug transcription factors are known to control EMT by downregulating cell-cell adhesion molecules, instigating loss of cell polarity, while elevating proliferative and migratory capacities. EMT is thus often linked to increased plasticity and stem cell activation during tissue regeneration. Bridging Schwann cells of the mammalian PNS share morphological and functional similarities with bridging glial cells of the zebrafish CNS [1, 7]. Yet, the extent of molecular similarities between these pro-regenerative glia remains to be determined.

Here, we used a battery of *ctgfa*-based genetic tools 1) to define the transcriptional profiles of bridging glial cells in zebrafish, 2) to determine the early injury-responses that direct *ctgfa*-mediated bridging, and 3) to elucidate molecular mechanisms that enable glial bridging. To these ends, we generated transgenic *ctgfa*/*gfap* dual reporter line to isolate and deep sequence *ctgfa*^+^*gfap*^+^ cells. The transcriptional profiles of zebrafish bridging glia shared similarities with the Repair Schwann cells that bridge transected nerves in the mammalian PNS. We identified a Yap-Ctgf-EMT signaling axis that directs glial bridging and functional regeneration. Yap activation correlates with *ctgfa* expression and is required for bridging and functional SC repair. Regulation of *ctgfa* expression converges onto a 1Kb *cis*-regulatory element that directs *ctgfa*-dependent transcription following injury. Yap-Ctgf signaling upregulates *twist* and *zeb* transcription factors, which in turn promote partial EMT and support SC regeneration. Our findings unraveled the molecular identity of bridging glial cells in zebrafish and revealed a previously unrecognized gene regulatory network that directs glial bridging following SCI.

## RESULTS

### Molecular profiling of *ctgfa*^+^*gfap*^+^ bridging glial cells after spinal cord injury

We devised a Fluorescence Activated Cell Sorting and RNA deep sequencing (FACS-seq) strategy to profile the transcriptome of bridging glial cells using Ctgfa and Gfap (Fig. 1). Ctgfa expression was previously shown to demarcate bridging glia at the lesion core and their predicted progenitors in the ventral ependyma proximal to the lesion [1]. To specifically label bridging glia, we generated transgenic *-5.5Kb-ctgfa:*mCherry reporter line in combination with the previously established *gfap:*EGFP transgene (mi2002, [11]) (Fig 1A). Broadly expressed in developing zebrafish animals, *ctgfa*-driven mCherry expression co-localized with *gfap*:EGFP in the ventral floor plate at 5 days post-fertilization (Fig. S1A). Consistent with its injury-induced expression, mCherry expression was not detectable in unlesioned adult SCs but was upregulated within SC tissue after injury (Fig. S1B). Longitudinal SC sections from adult *ctgfa:*mCherry;*gfap:*EGFP animals confirmed mCherry and EGFP expression in bridging glia as early as 5 days post-injury (dpi) (Fig. 1 A, B). mCherry expression was also broadly induced in *gfap*^+^ ependymal progenitors at 1 wpi (Fig. S1B), and localized to ventral ependymal cells at 10 dpi (Fig. 1 A, B). These results recapitulated the expression patterns of endogenous *ctgfa* transcripts and of a previously established -5.5Kb-*ctgfa*:EGFP reporter line after SCI [1]. Thus, we used the *ctgfa/gfap* dual reporter line to isolate *ctgfa*^+^*gfap*^+^ glial cells.

**Figure 1.**
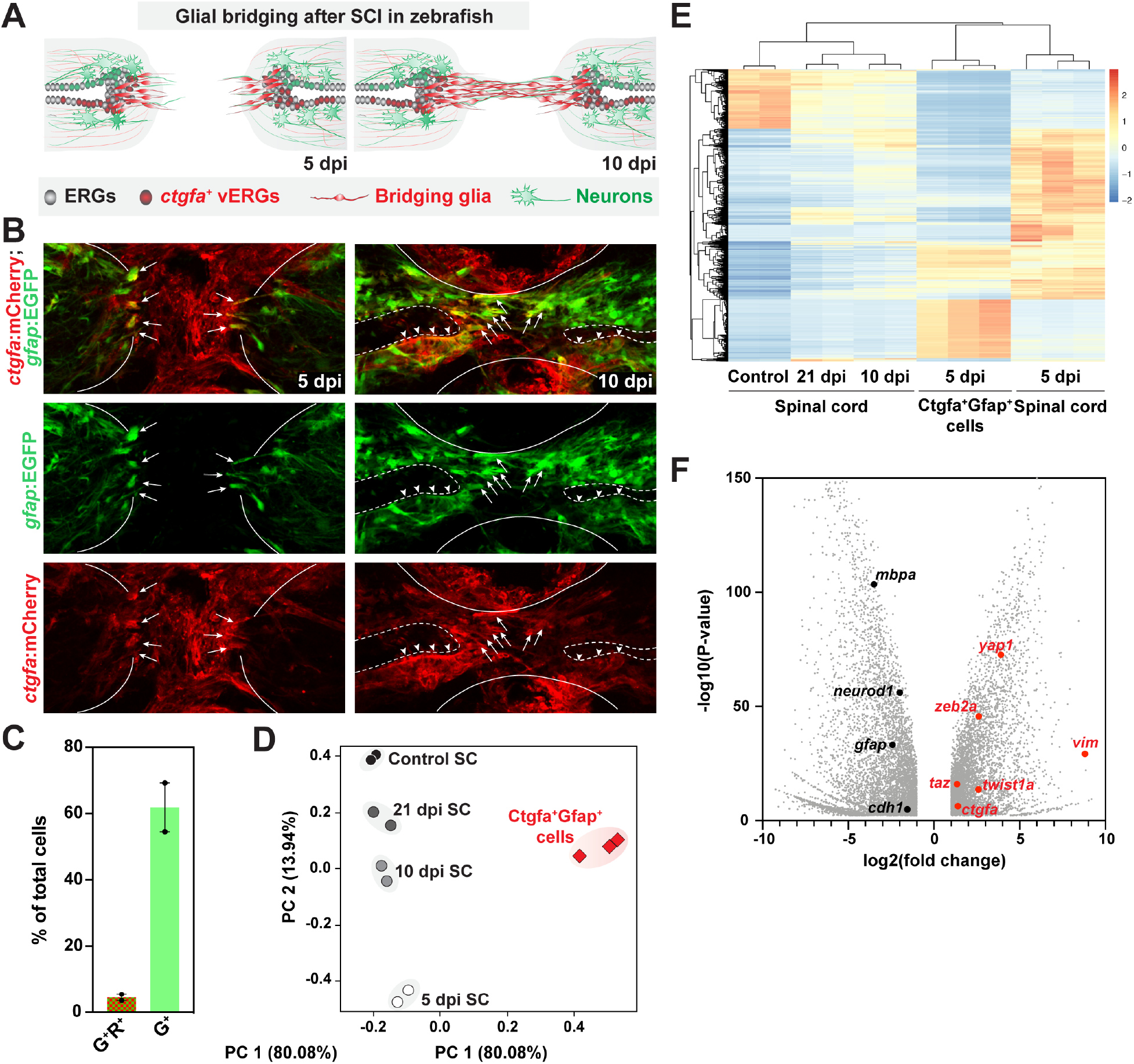
Molecular Profiling of bridging glial cells in zebrafish. **(A)** Schematic representation of zebrafish SCs at 5 and 10 dpi. Ependymal radial-glial (ERG) progenitors profilerate rostral and caudal to the lesion. The central canal expands proximal to the lesion from the rostral and caudal ends. *ctgfa* is expressed in a subset of ERGs at 5 dpi (red), and localizes to ventral ERGs (vERGs) at 10 dpi. Early bridging glia (red) emerge at 5 dpi and a glial bridge is formed by 10 dpi. **(B)** mCherry and EGFP immunostaining in *ctgfa*:mCherry;*gfap*:EGFP dual reporter line. Shown are longitudinal SC sections from adult animals at 5 dpi. Lines delineate the outer edges of the SC. Dashed lines outline the central canal. Arrows point to mCherry/EGFP double positive bridging glia. Ventral ERGs proximal to the lesion (arrowheads) are also positive for mCherry/EGFP expression. **(C,D)** Capture and deep RNA sequencing of bridging glia. Ctgfa^+^Gfap^+^ cells were sorted from *ctgfa*:mCherry;*gfap*:EGFP reporter animals at 5 dpi. Each replicate represents cells that were sorted from 40-50 SCs. Bulk SC tissues from 5, 10, and 21 dpi, as well as uninjured control SCs were deep sequenced. For bulk sequencing, 10 SCs from wild-type animals were pooled for each replicate and time point. Two biological replicates were used for control, 10 dpi, and 21 dpi SCs. Three biological replicates were used at 5 dpi. mCherry^+^EGFP^+^ (R^+^G^+^) and EGFP^+^ (G^+^) cells comprised 3% and 60% of total dissociated cells, respectively (C). PCA scatter plot of gene expression shows the variances between biological replicates (D). The percentages on each axis represent the percentage of variation explained by the principal components. **(E)** Heat map representation and hierarchical clustering of genes that are differentially expressed between groups. Color scale in the dendrogram represents relative expression levels: blue indicates down regulated genes, red indicates up-regulated genes, and yellow represents unchanged gene expression levels. **(F)** Volcano plot representation of genes that are significantly enriched in sorted Ctgfa^+^Gfap^+^ cells relative to control SCs. Genes with log2 (fold change) between −1 and 1, or with –log10 (P-value) less than 1 were considered insignificant. *ctgfa*, *yap1*, *taz*, *twist1a*, *zeb2a*, and *vim* are enriched in sorted cells. *gfap*, *neurod1*, *mbpa*, and *cdh1* transcript levels are attenuated in sorted cells relative to control SCs.

To define the molecular signature of pro-regenerative bridging glia, we performed FACS-seq on *ctgfa*^+^*gfap*^+^ cells and bulk SC tissue (Fig. 1 C-E). Complete SC transection was performed on *ctgfa/gfap* dual reporter animals. Single reporters and wild type animals were used as controls. At 5 dpi, 2 mm SC tissue samples including the lesion site were harvested and dissociated for FACS sorting. As a negative control, we collected SC tissue from sham injured animals (data not shown). From these experiments, we were able to collect mCherry^+^EGFP^+^ cells from animals with transected SCs but not from sham-injured animals. These results confirmed the injury-induced nature of Ctgfa and the emergence of *ctgfa*^+^*gfap*^+^ cells after injury. mCherry^+^EGFP^+^ cells comprised ~3% of the total dissociated cells at 5 dpi, while ~60% of the cells showing exclusive EGFP expression (Fig. 1 C). RNA samples from isolated *ctgfa*^+^*gfap*^+^ cells and bulk SC tissue at 5 dpi were deep sequenced. SC tissue samples from 10 and 21 dpi as well as uninjured control samples were also sequenced (Fig. 1 D, E). Principle component and heat map analyses revealed clustering of biological replicates, and highlighted the extent of molecular regeneration between 5, 10 and 21 dpi relative to control samples.

We next performed differential gene expression analysis to explore molecular mechanisms of glial bridging (Fig. 1 E). As predicted by sorting mCherry^+^EGFP^+^ cells, our FACS-seq approach enriched for *ctgfa* expression (Fig. 1 E). *ctgfa* is expressed in multiple cell types around the lesioned SC; yet, its expression is primarily confined to bridging glia and ventral ependymal progenitors within dissected SC tissue. We thus observed elevated *ctgfa* expression in *ctgfa*^+^*gfap*^+^ cells relative to bulk SC tissue. Conversely, *gfap* is expressed in ~60% of the total cells dissociated from SC tissue, and Ctgfa^+^ cells constitute only a minor subset of Gfap expressing cells. Consequently, *gfap* expression was attenuated in isolated *ctgfa^+^gfap^+^* cells relative to bulk SC tissue. We also observed that neuronal and oligodendrocyte markers, such as *neurod1* and *mbpb*, were depleted in isolated *ctgfa*^+^*gfap*^+^ cells, suggesting that isolated *ctgfa*^+^*gfap*^+^ cells were depleted of neurons and oligodendrocytes. These studies generated the first transcriptional blueprints for pro-regenerative bridging glia in adult zebrafish.

### Yap is activated in Ctgfa-expressing cells after spinal cord injury

The Hippo signaling pathway and its downstream co-activators Yap and Taz play important roles in injury response and tissue regeneration in multiple tissue contexts. Similar to *ctgfa* expression after SCI, the Yap target genes, *cyr61* and *gli2a*, were upregulated in bulk SC tissue and enriched in isolated *ctgfa*^+^*gfap*^+^ cells at 5 dpi relative to uninjured controls (Fig. 2 A). Moreover, *yap1*, *taz*, and their transcription factor effectors *tead1a* and *tead1b* were upregulated in injured SC tissue (Fig. 2A). However, while *taz* was ~2-fold enriched in isolated *ctgfa*^+^*gfap*^+^ cells, *yap1* and the *tead* transcription factors were only mildly enriched by FACS-seq, reflecting their ubiquitous expression pattern across multiple SC cell types. These gene expression studies suggested that Yap signaling is activated upstream of Ctgfa during glial bridging.

**Figure 2.**
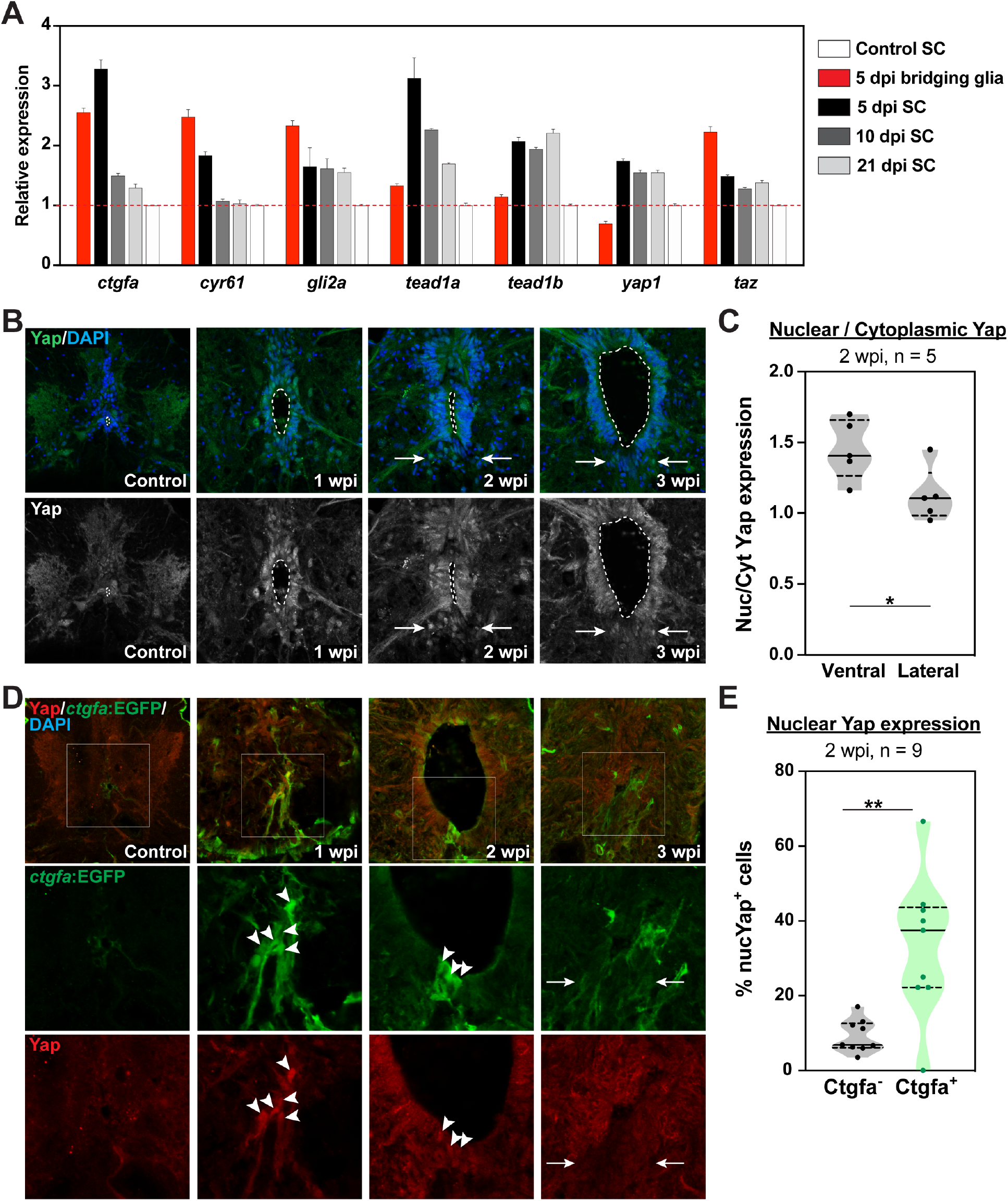
Localized activation of Yap signaling in ERG progenitors after SCI. **(A)** Expression of Yap related genes by FACS-seq. Relative expression represents the average fpkm for each gene relative to its expression in control SCs. In addition to *ctgfa*, the Yap target genes *cyr61* and *gli2a* are upregulated in bulk SC tissue and enriched in sorted Ctgfa^+^Gfap^+^ cells after SCI. **(B, C)** Yap expression at 1, 2, and 3 wpi, and in uninjured control tissue. Yap immunostaining was performed on SC cross sections from wild-type zebrafish. Dotted lines delineate the edges of the central canals. Yap showed broad nuclear expression in ERG progenitors at 1 wpi. Nuclear Yap expression is preferentially localized to the ventral ependymal domains at 2 and 3 wpi (arrows). For quantification, nuclear and cytoplasmic Yap expression were quantified in ventral and lateral ependymal domains at 2 wpi. Nuclear-to-cytoplasmic ratios were averaged for 2 sections per animal. Five animals from 2 independent experiments were used. **(D, E)** Nuclear Yap expression correlates with Ctgf expression after SCI. Yap and EGFP immunostaining from *ctgfa*:EGFP reporter fish. Shown are SC cross sections at 1, 2, and 3 wpi, and in uninjured control tissue. Single channel views of *ctgfa*:EGFP (green) and Yap (red) are shown in high-magnification. Arrowheads point to individual cells that are positive for nuclear Yap and EGFP in ventral ependymal progenitors at 1 and 2 wpi. Arrows point to nuclear Yap and EGFP expression in the ventral ependyma at 3 wpi. For quantification, the number of nuclear Yap expressing cells were calculated within Ctgfa^-^ and Ctgfa^+^ expression domains at 2 wpi. Percent nuclear Yap expression relative to the total number of cells within each domain were averaged for 2 sections per animals. Nine animals from two independent experiments were used. \**P*<0.05 and \*\**P*<0.01.

To further examine Yap and Taz activation after SCI, we performed immunohistochemistry on on wild-type SCs at 1, 2, and 3 wpi as well as uninjured controls (Fig. 2 B, S2, and S3). Consistent with FACS-seq and RNA-seq data, Yap was broadly expressed in control and lesioned SCs between 1 and 3 wpi. Yet, the sub-cellular localization of Yap revealed dynamic changes during regeneration (Fig. 2 B and S2). At 1 wpi, nuclear Yap was broadly expressed around the ependyma, suggesting global transcriptional activation early after SCI. Yap expression showed preferential nuclear localization within ventral ependymal progenitor relative to either dorsal or lateral cells at 2 wpi. Although Yap expression was broadly downregulated at 3 wpi, Yap maintained its localization to the nuclei of ventral ERGs at this time point. Yap quantification in the nuclear and cytoplasmic compartments confirmed its nuclear enrichment (~1.4 fold nuclear to cytoplasmic expression) in ventral progenitors at 2 wpi (Fig 2 C). Yap expression was uniformly distributed between nuclei and cytoplasms in lateral progenitors at 2 wpi (~1 fold nuclear to cytoplasmic expression) (Fig. 2 C). By immunostaining, we also observed elevated Taz expression in injured SC sections relative to uninjured control SCs (Fig. S3). Similarly to Yap, Taz expression peaked between 1 and 2 wpi, and diminished toward baseline expression by 3 wpi. These experiments indicated that Yap and Taz are activated after SCI, and that the dynamics of Yap activation are analogous to *ctgfa* gene expression during glial bridging [1].

To examine whether Yap activation correlates with Ctgfa expression, we injured *-5.5Kb-ctgfa*:EGFP transgenic zebrafish [1] and performed time course analysis for Yap and EGFP expression (Fig. 2 D, E). Consistent with previously reported *ctgfa* expression, nuclear Yap colocalized with *ctgfa*-driven EGFP expression proximal to the lesion at 1 wpi. By 2 wpi, EGFP and nuclear Yap expression were more confined to ventral ependymal progenitors. At this time point, ~40% of total SC cells showed dual expression of nuclear Yap and *ctgfa*:EGFP, with ~5% of the cells expressing nuclear Yap in the absence of EGFPexpression (Fig. 2 E). Yap expression was decreased in ventral ependymal progenitor by 3 wpi, correlating with attenuated *ctgfa*:EGFP expression after glial bridge formation at later stages of regeneration (Fig. 2D). Together, these experiments indicated a correlation between nuclear Yap localization and Ctgfa expression during glial bridging.

### A 1Kb enhancer element directs *ctgfa* expression after SCI

Ctgfa is a target of Yap signaling in multiple tissue contexts, yet it remains unclear whether similar regulatory mechanisms applied during SC regeneration. We performed a comprehensive analysis of the *cis*-regulatory elements that direct injury-induced *ctgfa* expression in regenerating SC tissue (Fig. 3 A). Using *-5.5Kb-ctgfa*:EGFP, we previously showed that EGFP fluorescence recapitulates endogenous mRNA expression in ventral ependymal progenitors and in bridging glia during SC regeneration [1]. Bioinformatics analysis of the 5.5 Kb genomic region revealed predicted binding sites for Tead transcription factors 3 to 4 Kb upsteam of *ctgfa* translational start site (Fig. S4). Stable transgenic lines expressing an EGFP cassette downstream of the -4Kb- or -3Kb genomic regions were generated (*-4kb-ctgfa*:EGFP and *-3kb-ctgfa*:EGFP, respectively) (Fig. 3A). Multiple independent lines were generated for each transgene to control for possible positional effects that may impact transgene expression. At 10 dpi, *ctgfa*-driven EGFP and Gfap expression were assessed 100 μm (proximal) and 500 μm (distal) rostral to the lesion (Fig. 3 B). Similar to *-5.5Kb*-*ctgfa*:EGFP, only *-4Kb-ctgfa*:EGFP transgenic lines showed EGFP expression at 10 dpi; while EGFP expression was not detectable in *-3Kb-ctgfa*:EGFP lines (Fig. 3 C, D). These results suggested that *cis*-regulatory elements located between 3 and 4 Kb upstream of the *ctgfa* translational start site direct *ctgfa* expression after SCI. To determine whether this putative 1Kb enhancer element is sufficient to promote EGFP expression after SCI, we generated *1Kb-ctgfa*:EGFP transgenic line and examined EGFP expression following injury. EGFP was not detectable in uninjured controls, but was comparable to *-5.5Kb-* and *-4Kb-ctgfa*:EGFP expression at 10 dpi (Fig. 3 C, D). This expression pattern was recapitulated in three independent lines of the *1Kb*-*ctgfa*:EGFP transgene (Fig. S5). These results revealed a 1Kb enhancer element that directs injury-induced *ctgfa* expression after SCI.

**Figure 3.**
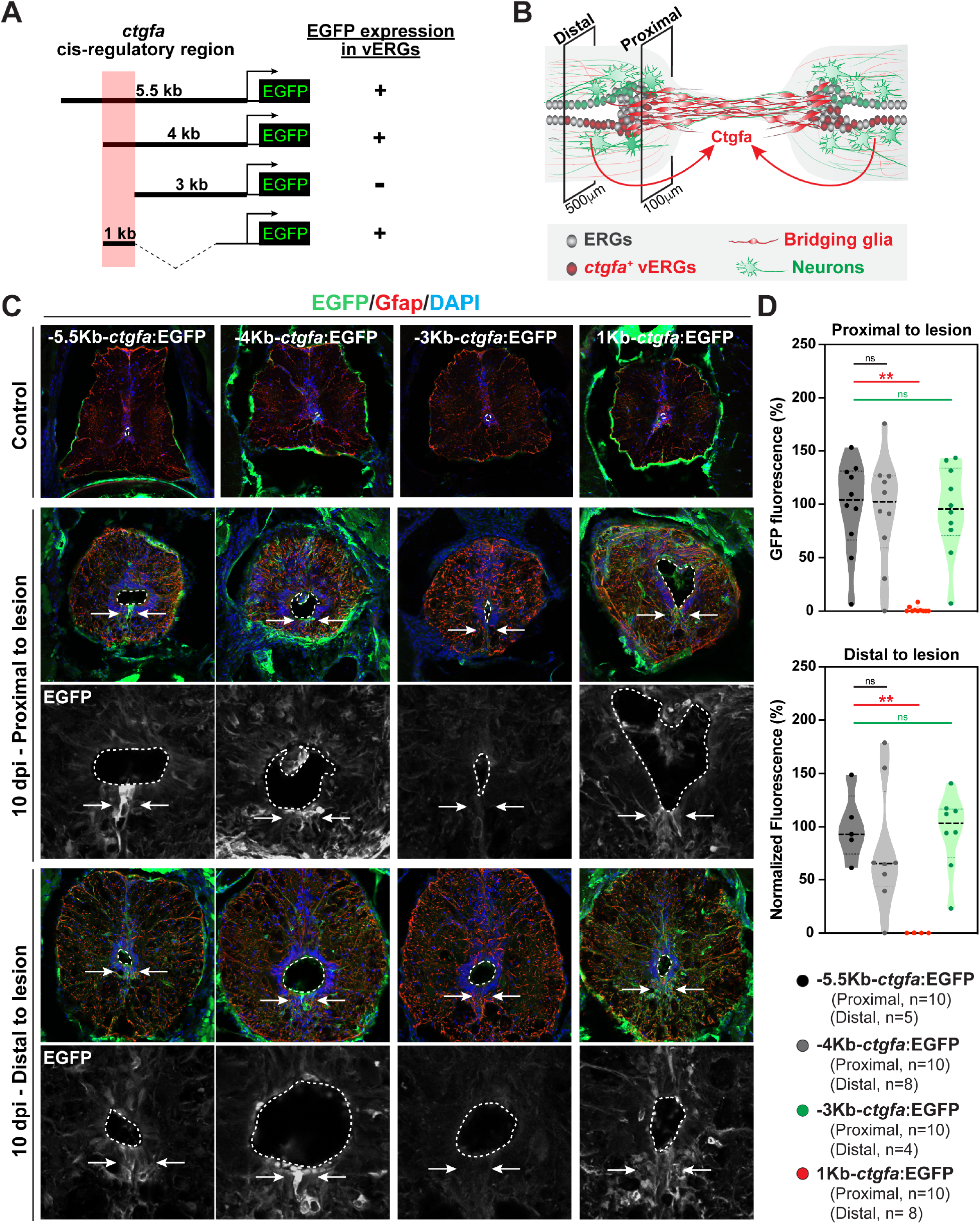
Regulation of *ctgfa* expression during SC regeneration. **(A)** A Yap-dependent enhancer element drives localized *ctgfa* expression after SCI. A series of transgene constructs were used to identify the *cis*-regulatory region that drives *ctgfa*-dependent EGFP expression after SCI. A minimum of 3 stable lines were generated for each transgene. A 1Kb enhancer element located 3-4 Kb from the translational start site of *ctgfa* was sufficient to recapitulate expression in ventral ependymal progenitors after injury. **(B)** Schematic of regenerating SC tissue in longitudinal view at 10 dpi. SC anatomy and gene expression show significant differences along the rostro-caudal axis of injured SCs. Shown are the proximal and distal SC levels that were analyzed in this experiment. Proximal and distal tissue sections are located 100 and 500 μm rostral to the lesion core, respectively. (**C, D**) EGFP and Gfap immunostaining assessed reporter expression at 10 dpi and in uninjured tissue. Cross sections are shown at the proximal and distal levels schematized in panel B. SC tissues from *-5.5Kb-ctgfa*:EGFP, *-4Kb-ctgfa*:EGFP, *-3Kb-ctgfa*:EGFP, and *1Kb-ctgfa*:EGFP transgenic animals are shown. Dotted lines delineate central canal edges. Arrows point to EGFP expression in ventral ependymal progenitors. EGFP expression in *-4Kb-ctgfa*:EGFP and *1Kb-ctgfa*:EGFP lines recapitulated *-5.5Kb-ctgfa*:EGFP expression. EGFP expression is not detectable in *-3Kb-ctgfa*:EGFP animals after SCI. EGFP fluorescence was quantified at the proximal and distal levels. Ten animals from three independent experiments were analyzed at the proximal level. 4-8 animals from two independent experiments were analyzed at the distal level. \*\**P*<0.01; ns, not significant.

### Yap is required for Ctgfa expression and for functional SC repair

To determine the effects of Yap inactivation, we expressed dominant negative Yap under control of a heat-inducible promoter (*hsp70*:dsRed-dnYap) (Fig. 4). We first combined dnYap expressing and *-5.5Kb-ctgfa*:EGFP transgenes to examine *ctgfa*-driven EGFP expression upon Yap inactivation. *hsp70*:dsRed-dnYap;*ctgfa*:EGFP (Tg^+^) zebrafish and *ctgfa*:EGFP (Tg^-^) controls were subjected to SC transections followed by daily heat shocks to induce dnYap expression. EGFP expression was observed in ventral ependymal progenitors of *ctgfa*:EGFP controls at 10 dpi, but was attenuated in dnYAP-expressing animals (Fig. 4 A). Quantification showed a significant decrease in EGFP expression in the ventral SCs of dnYap-expressing animals relative to their control siblings (Fig. 4 B). These results indicated that Yap promotes *ctgfa* expression in ventral ependymal progenitors after SCI.

**Figure 4.**
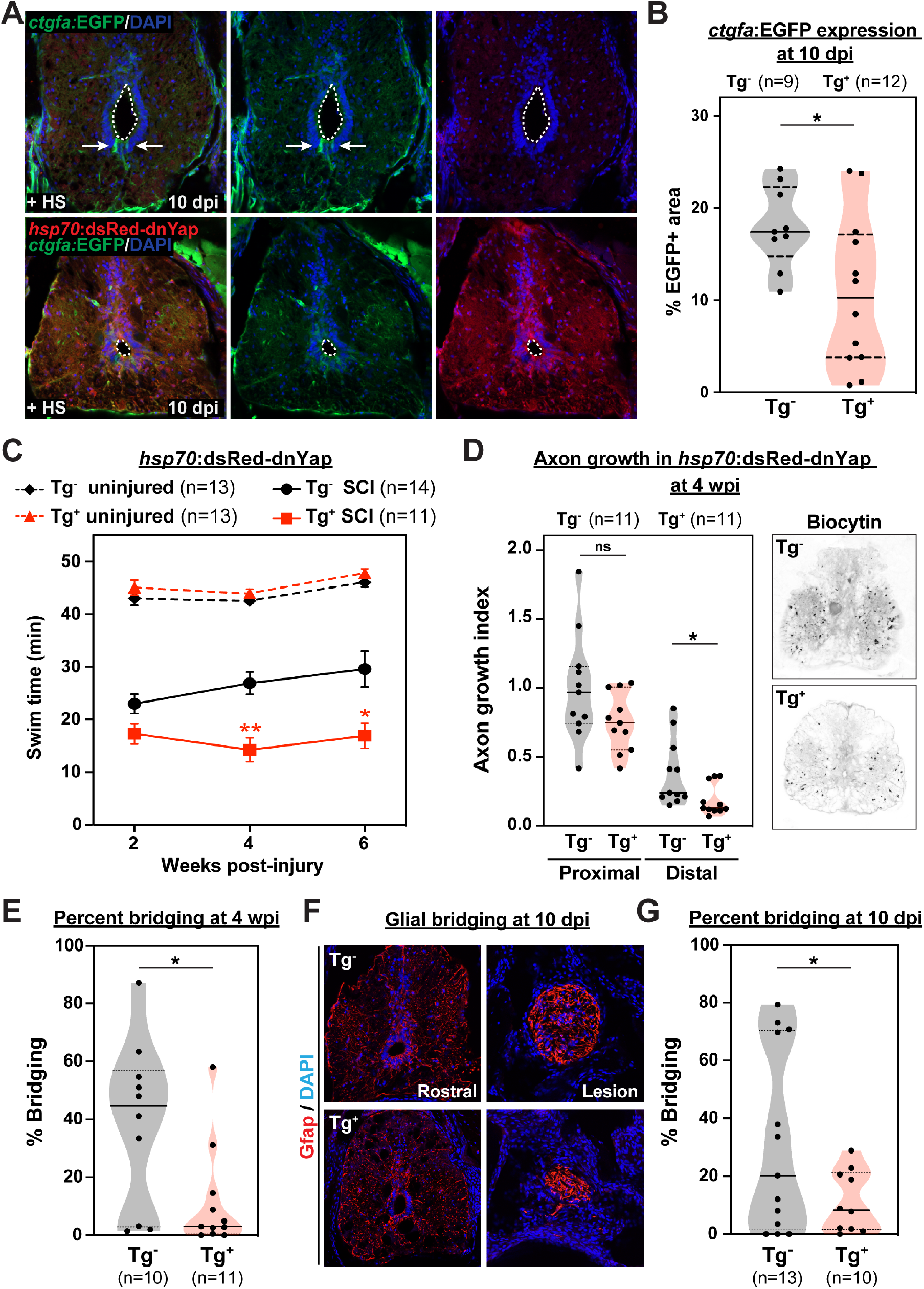
Yap promotes *ctgfa* expression, glial bridging, and SC regeneration. **(A, B)** EGFP immunostaining in *ctgfa*:EGFP;*hsp70*:dsRed-dnYap (Tg^+^) SCs at 10 dpi. *ctgfa*:EGFP (Tg^-^) siblings were used as controls. Tg+ and Tg-animals were subjected to SCI and daily heat shocks (+HS) to induce dnYap transgene expression. EGFP is expressed in ventral ependymal progenitors at 10 dpi (arrows) in control SCs, but is attenuated in dnYap expressing SCs. Dotted lines delineate central canal edges. For quantification, the area of EGFP fluorescence was calculated for 2 sections per animals. Nine Tg^-^ and 12 Tg^+^ animals from two independent experiments were used. **(C)** Swim assays determined motor function recovery of 11 *hsp70*:dsRed-dnYap (Tg^+^, SCI, red) and 14 wild-type (Tg^-^, SCI, black) siblings at 2, 4, and 6 wpi. For controls, 13 dnYap overexpressing (Tg^+^, uninjured, dashed red) and 13 wild-type (Tg^-^, uninjured, dashed black) animals were analyzed. Experimental and control groups were subjected to daily heat shocks. Statistical analyses of swim times are shown for injured dnYap relative to injured wild-type siblings. **(D)** Anterograde axon tracing in dnYap-expressing zebrafish at 4 wpi. Biocytin axon tracer was applied rostrally and analyzed at 100 μm (proximal) and 500 μm (distal) caudal to the lesion. Representative traces of biocytin are shown for Tg^+^ and Tg^-^ animals at the proximal level. Quantification represents 11 dnYap overexpressing and 11 wild-type zebrafish from two independent experiments. **(E)** Glial bridging in dnYap-expressing zebrafish at 4 wpi. Gfap immunostaining was used to quantify glial bridging at 4 wpi in 11 dnYap-expressing and 10 wild-type zebrafish. **(F, G)** Glial bridging in dnYap-expressing zebrafish at 10 dpi. Representative immunohistochemistry shows the Gfap^+^ bridge at the lesion site relative to the intact SC rostral to the lesion in dnYap expressing (Tg^+^) and control siblings (Tg^-^). Percent bridging is quantified for 10 dnYap-expressing and 13 wild-type zebrafish from two independent experiments. \**P*<0.05; ns, not significant.

We then assessed the outcomes of Yap inactivation on functional and anatomical SC repair (Fig. 4 C). Swim assays revealed impaired functional recovery in dnYap-expressing zebrafish given daily heat shocks and assessed at 2, 4, and 6 wpi. Swim capacity was comparable between uninjured dnYap-expressing and wild-type siblings after 2, 4 and 6 weeks of daily heat shocks, indicating contextual effects of dnYap expression upon injury. Anterograde axon tracing with Biocytin indicated that axon regeneration across the lesion site was significantly reduced distal to the lesion site at 4 wpi (Fig. 4 D). At this time point, quantification of glial bridge formation revealed ~75% less bridging in dnYap expressing animals relative to transgene negative siblings (Fig. 4 E). We next examined glial bridging at earlier bridging stages to assess glial bridging defects upon dnYap expression (Fig. 4 F, G). At 10 dpi, Gfap immunostaining revealed the formation of an organized circular bridge at the lesion core. However, the glial bridges formed upon dnYap expression were 50% reduced and irregularly shaped relative to controls. These results revealed that Yap promotes glial bridging and functional SC repair.

### Localized EMT directs glial bridging downstream of Ctgfa

Ctgf proteins exert diverse functions by binding various cell surface receptors, growth factors, and extracellular matrix proteins [12]. By FACS-seq, *twist1a* and *zeb2a* transcription factors are upregulated in ventral ependymal progenitors at 5 dpi (Fig. 5 A). *in situ* hybridization confirmed that *twist1a* transcripts showed elevated expression in ventral ependymal progenitors at 2 wpi, (Fig. 6 B). Localized treatment with human recombinant C-terminal CTGF (CTGF-CT), which is sufficient to promote glial bridging and functional regeneration [1], induces ectopic Twist expression in ependymal cells at 2 wpi (Fig. 6 C). These experiments indicated that the transcription factor *twist1a* is a downstream effector of *ctgfa* during glial bridging, and suggested that *twist* and *zeb* transcription factors drive EMT downstream of Yap-Ctgf signaling after SCI.

**Figure 5.**
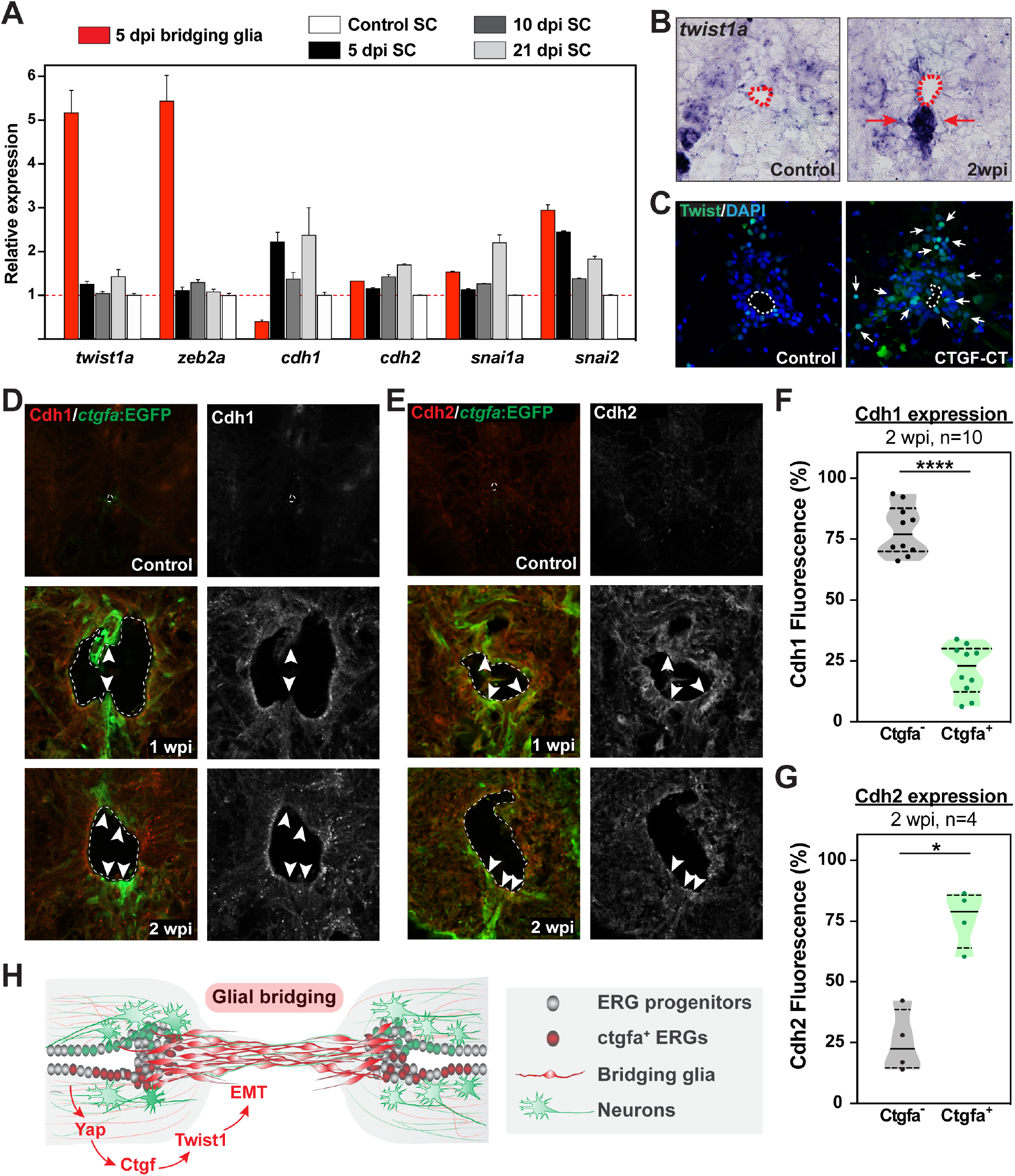
Localized EMT directs glial bridging downstream of Ctgfa. **(A)** Expression of EMT related genes by FACS-seq. Relative expression represents the average fpkm for each gene relative to its expression in uninjured control SCs. *twist1a* and *zeb2a* are ~5-fold enriched in sorted Ctgfa^+^Gfap^+^ cells. *snai1a*, and *snai2* are upregulated in bulk SCs and enriched in sorted cells. *cdh1* is upregulated in bulk SCs after SCI, but is depleted in sorted cells*. cdh2* expression did not show significant changes by FACS-seq. **(B)** *twist1a in situ* hybridization in wild-type SCs at 1, 2, and 3 wpi, and in uninjured controls. SC cross sections proximal to the lesion from the rostral side are shown. Dashed lines delineate central canal edges. Arrows point to *twist1a* expression in ventral ependymal progenitors. **(C)** Twist immunostaining in wild-type zebrafish treated with human recombinant C-terminal CTGF (CTGF-CT). Vehicle-treated siblings were used as controls. At 2 wpi, Twist1 is detected in few ventral ependymal progenitors in vehicle treated controls, but showed ectopic expression upon CTGF-CT treatment. **(D)** Cdh1 and EGFP immunostaining in *ctgfa*:EGFP zebrafish at 1 wpi, 2 wpi, and in uninjured controls. SC cross sections are shown. Dotted lines delineate central canal edges. Arrowheads points to domains of diminished Cdh1 expression and increased EGFP expression. **(E)** Cdh2 and EGFP immunostaining in *ctgfa*:EGFP zebrafish at 1 wpi, 2 wpi, and in uninjured controls. SC cross sections are shown. Dotted lines delineate central canal edges. Arrowheads points to domains of co-expression of Cdh2 and EGFP. **(F, G)** Quantification of Cdh1 (F) and Cdh2 (G) expression in Ctgfa^-^ and Ctgfa^+^ cells at 2 wpi. Ten animals and 4 animals were used for Cdh1 and Cdh2 quantification, respectively. \**P*<0.05 and \*\*\*\**P*<0.0001. **(H)** Schematic model for localized activation of ependymal progenitors by Yap-Ctgf-EMT signaling to promote glial bridging and SC regeneration in zebrafish.

*twist1* and *zeb2* are key transcription factor effectors of EMT. Hallmarks of EMT include concomitant downregulation of epithelial E-Cadherin (Cdh1) and upregulation of N-cadherin (Cdh2). We found that *cdh1* is depleted in isolated *ctgfa*^+^*gfap*^+^ cells at 5 dpi despite its upregulation in bulk SC tissue after SCI (Fig. 5 A). *cdh2* is expressed isolated *ctgfa*^+^*gfap*^+^ cells, suggesting these cells are undergoing a transition into mesenchymal cell fate. Additional mesenchymal markers, *snai1a* and *snai2*, are upregulated in injured SC tissue and enriched in bridging glia at 5 dpi. By immunohistochemistry, Cdh1 was specifically downregulated in *ctgfa*^+^ cells at 1 and 2 wpi (Fig. 5 D). Cdh1 expression was increased in lateral ERGs at these time points, and broadly attenuated toward baseline expression by 3 wpi. On the other hand, Cdh1 and Vim showed dynamic gene expression changes after SCI. High levels of Cdh1 expression were detectable in *ctgfa*^+^ cells at 1 and 2 wpi. Vimentin was globally upregulated at 1 wpi, showed preferential expression in ventral ependymal progenitors at 2 wpi, and localized to the progenitor motor domain by 3 wpi (Fig. S6). These results are consistent with the FACS-seq data and suggested that *ctgfa*^+^ cells undergo a mesenchymal transition during glial bridging.

## DISCUSSION

This study defines the transcriptional profiles of bridging glial cells in zebrafish, and found that Yap-Ctgf-EMT signaling axis mediates localized injury-responses and directs glial bridging after SCI. Using newly generated *ctgfa*/*gfap* dual reporter line, we isolated and deep sequenced *ctgfa*^+^*gfap*^+^ cells. Yap nuclear localization, Ctgf expression, and EMT markers impinge on ventral ependymal progenitors, instructing a glial bridging cell fate that supports innate regeneration in zebrafish. We show that Yap promotes Ctgfa expression, glial bridging and functional SC repair, and that Ctgfa regulation is driven by a 1Kb regeneration enhancer that drives its expression after SCI. Yap-Ctgf signaling activates Twist driven EMT during glial bridging and SC regeneration.

Glial cell responses are thought to dictate SCI outcomes across species. Following SC transection in zebrafish, specialized glial cells connect the severed SC ends, and axons regrow along these bridging glia [1]. From a translational standpoint, we envision shifting mammalian glia towards a bridging phenotype to abate scarring and support axon regrowth. Practically, this outcome requires detailed understanding of the cells and molecules that induce glial bridge formation. Our study uncovered molecular signature of bridging glial cells in zebrafish. We found that glial bridging shares morphological and molecular hallmarks with Schwann cell mediated bridging after peripheral nerve injuries in mammals. Similar to activated Schwann cells, bridging glia emerge via bidirectional cell migration from the rostral and caudal spinal stumps. Peripheral nerve injury induces dynamic changes in Yap and Taz expression within the spinal cord [17].While hallmarks of Vegf signaling activity are present in bridging glia by FACS-seq analysis, the mechanisms that direct proper migration and alignment of bridging glia along the rostro-caudal axis of the lesioned SC remain to be determined.

Yap signaling plays important roles in CNS development and tumorigenesis. This pathway is regulated by various upstream regulators, including cell polarity, adhesion proteins, Wnt/β-catenin and MAPK pathways. Of high relevance to tissue injury and regeneration, Yap is a mechanoresponsive signaling pathway that relays cellular stress into transcriptional responses [13]. One of Yap’s mechanosensing functions is to survey the levels of filamentous F-actin, and manipulating F-actin levels is sufficient to shuttle Yap into or out of the nucleus [14]. Active Yap signaling maintains stemness in various stem cell types, including neural and glial progenitors [15]. Yap inactivation promotes stem cell differentiation, while Yap activation is sufficient to revert differentiated cells back to a tissue-specific stem/progenitor cell state [16]. Complete SC transection in zebrafish results in pronounced mechanical stress, manifested by prolonged expansion of the central canal proximal to the lesion from the rostral and caudal ends. Our results are consistent with a model in which Yap mechanosenses this stress, resulting in activation of ependymal progenitor cells. Why and how Yap activation confines to specific niches of progenitors warrant further investigation.

Our studies shed light on glial bridging as an effective, natural mechanism of SC repair. Equipped with a superior regenerative capacity relative to the mammalian CNS, zebrafish SCs trigger pro-regenerative responses that include glial cell activation and EMT-driven bridging following injury. We expect further investigation into bridging glial cell fate to springboard translational applications to improve bridging and regeneration in the mammalian CNS.

---

**Figure S1.**
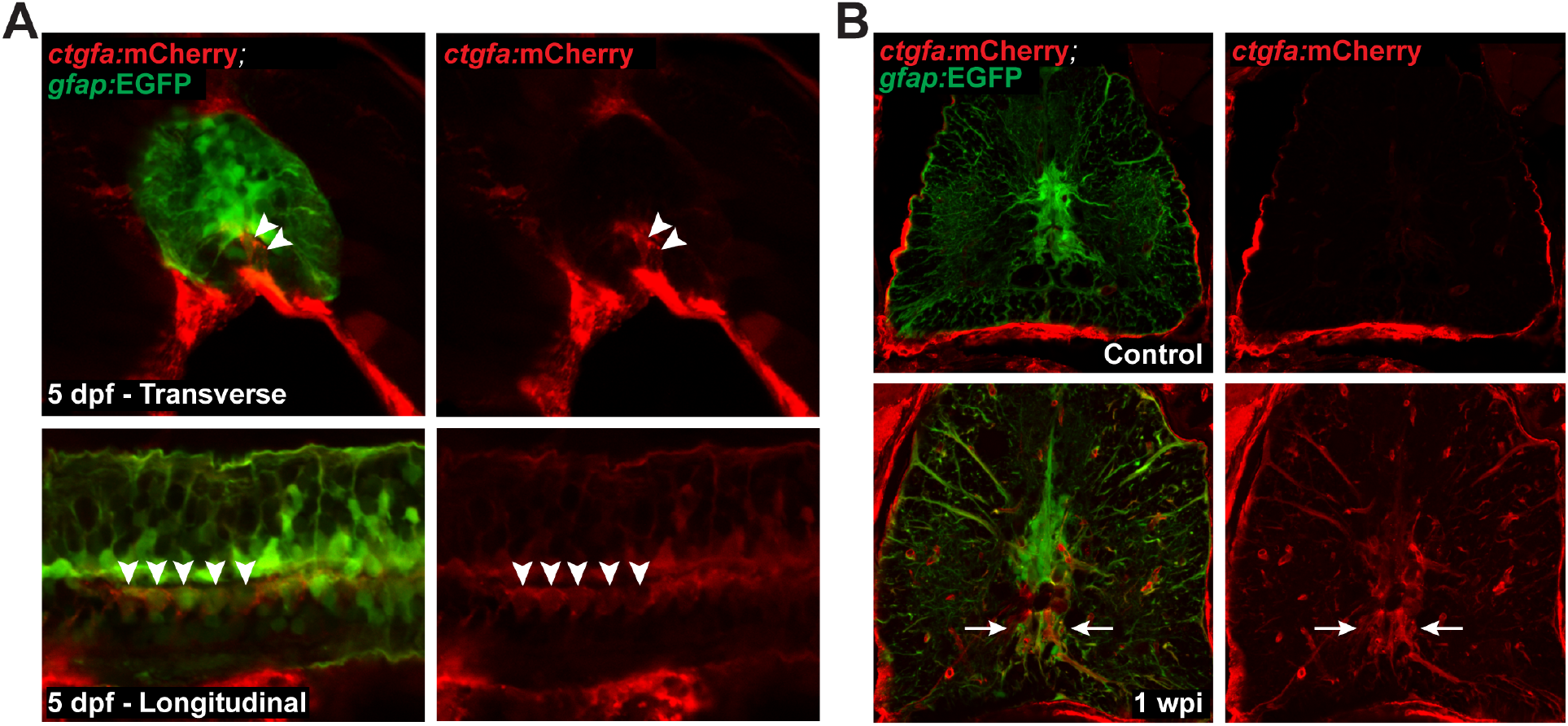
Generation of *ctgfa*:mCherry reporter line. **(A)** mCherry and EGFP expression in *ctgfa*:mCherry;*gfap*:EGFP dual reporter animals at 5 days post-fertilization (dpf). Transverse and longitudinal SC sections are shown. Arrowheads point to mCherry/EGFP double positive cells in the floor plate of the developing SC. **(B)** mCherry and EGFP expression in adult *ctgfa*:mCherry;*gfap*:EGFP dual reporter animals. Transverse SC sections at 1 wpi and in uninjured controls are shown. mCherry expression is induced within SC tissues upon injury. Arrows point to ventral ependymal progenitors that are mCherry and EGFP positive.

**Figure S2.**
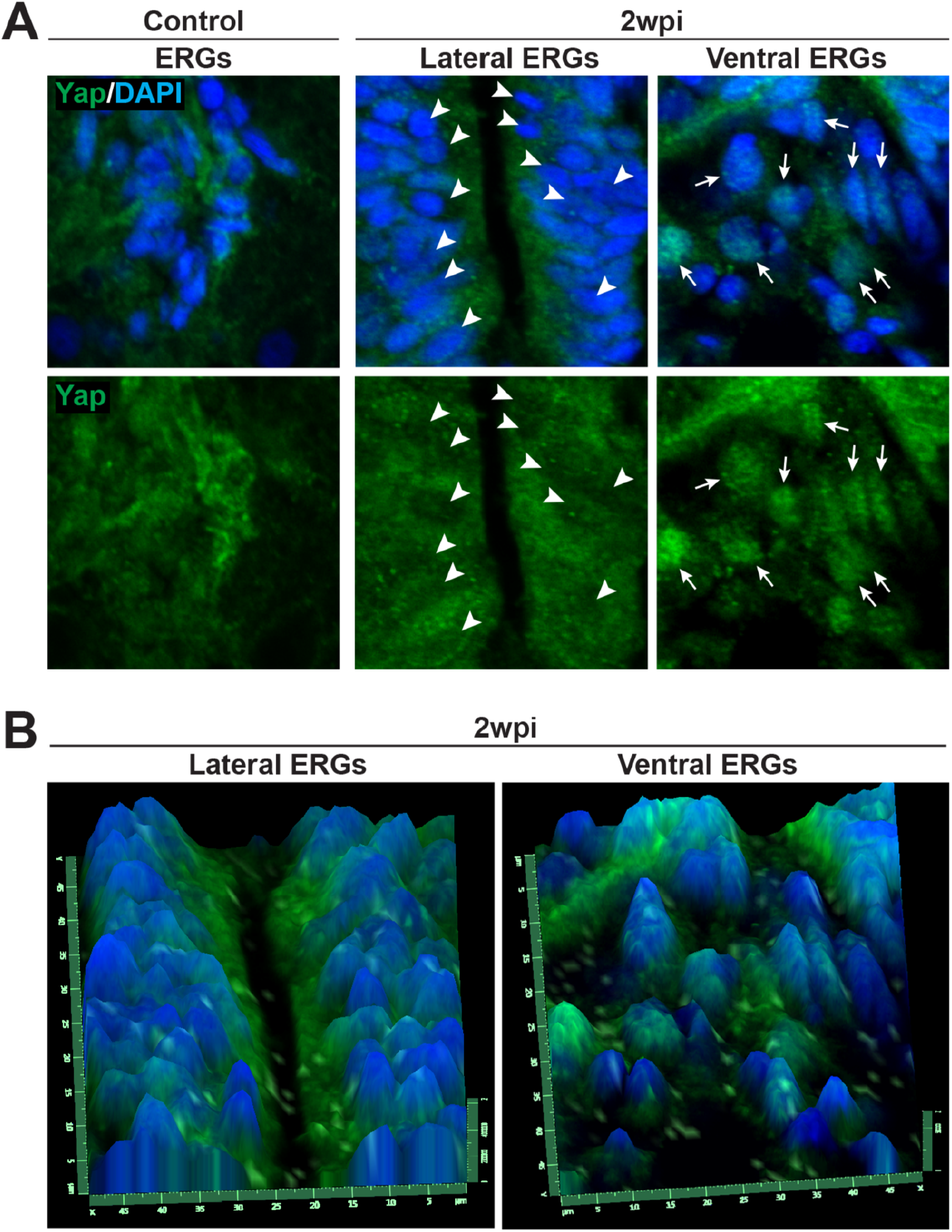
Yap expression in adult zebrafish after SCI. **(A)** High magnification views of Yap expression in wild-type zebrafish at 2 wpi and in uninjured controls. Yap immunostaining was performed on SC cross sections. All ependymal progenitors are shown in a single panel in control animals, which have constricted central canals. Lateral and ventral ependymal progenitors are shown in separate panels at 2 wpi. Arrowheads point to nuclei with attenuated Yap expression in lateral ERGs at 2 wpi. Arrows point to nuclei with elevated Yap expression in ventral ERGs at 2 wpi. **(B)** Three-dimensional reconstruction of Yap expression at 2 wpi.

**Figure S3.**
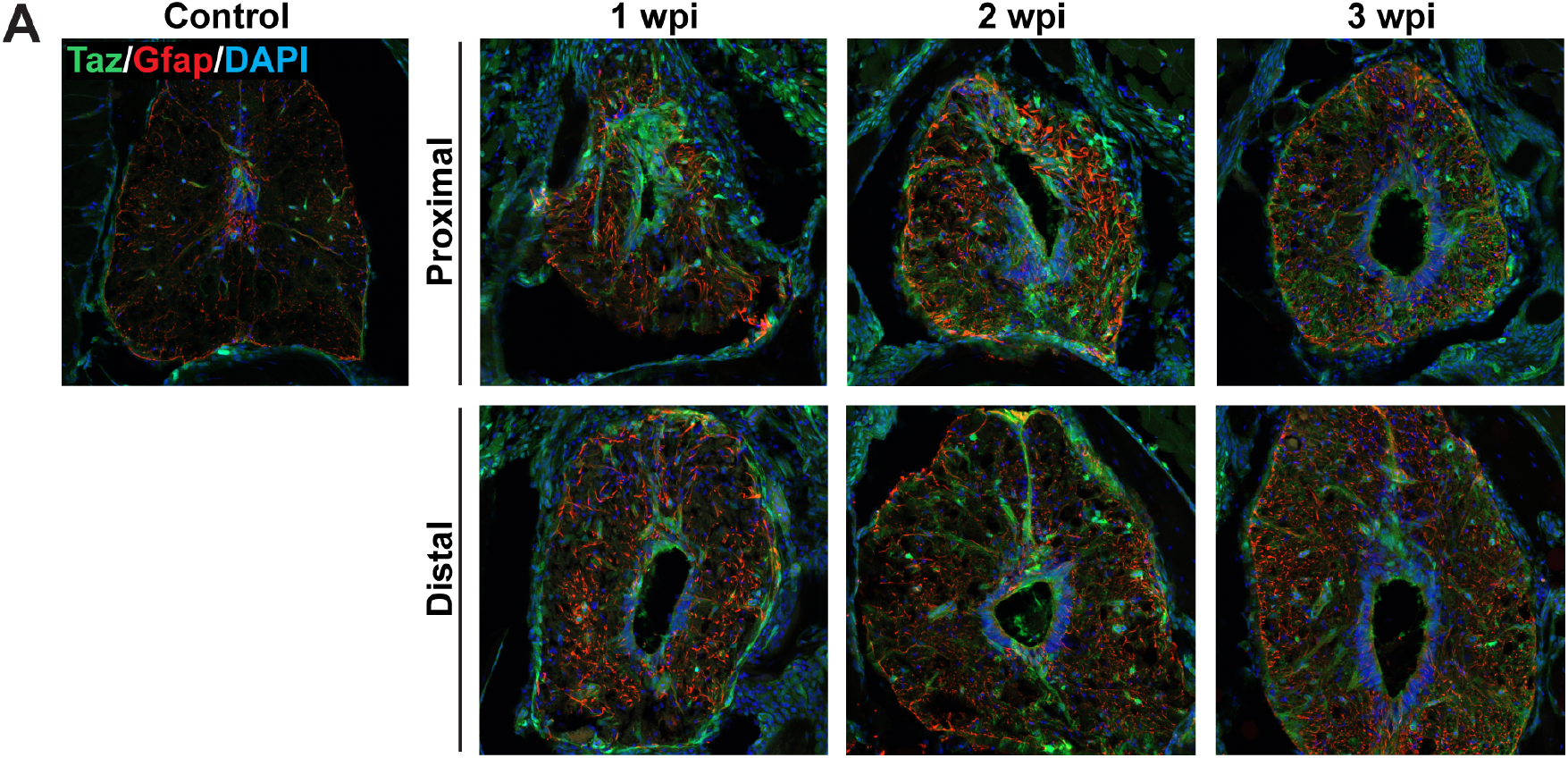
Taz expression in wild-type zebrafish after SCI. Taz and Gfap immunostaining was performed at 1, 2, and 3 wpi, and in uninjured control tissue. SC cross sections proximal and distal to the lesion site are shown. Taz expression is broadly upregulated in ependymal progenitors after SCI.

**Figure S4.**
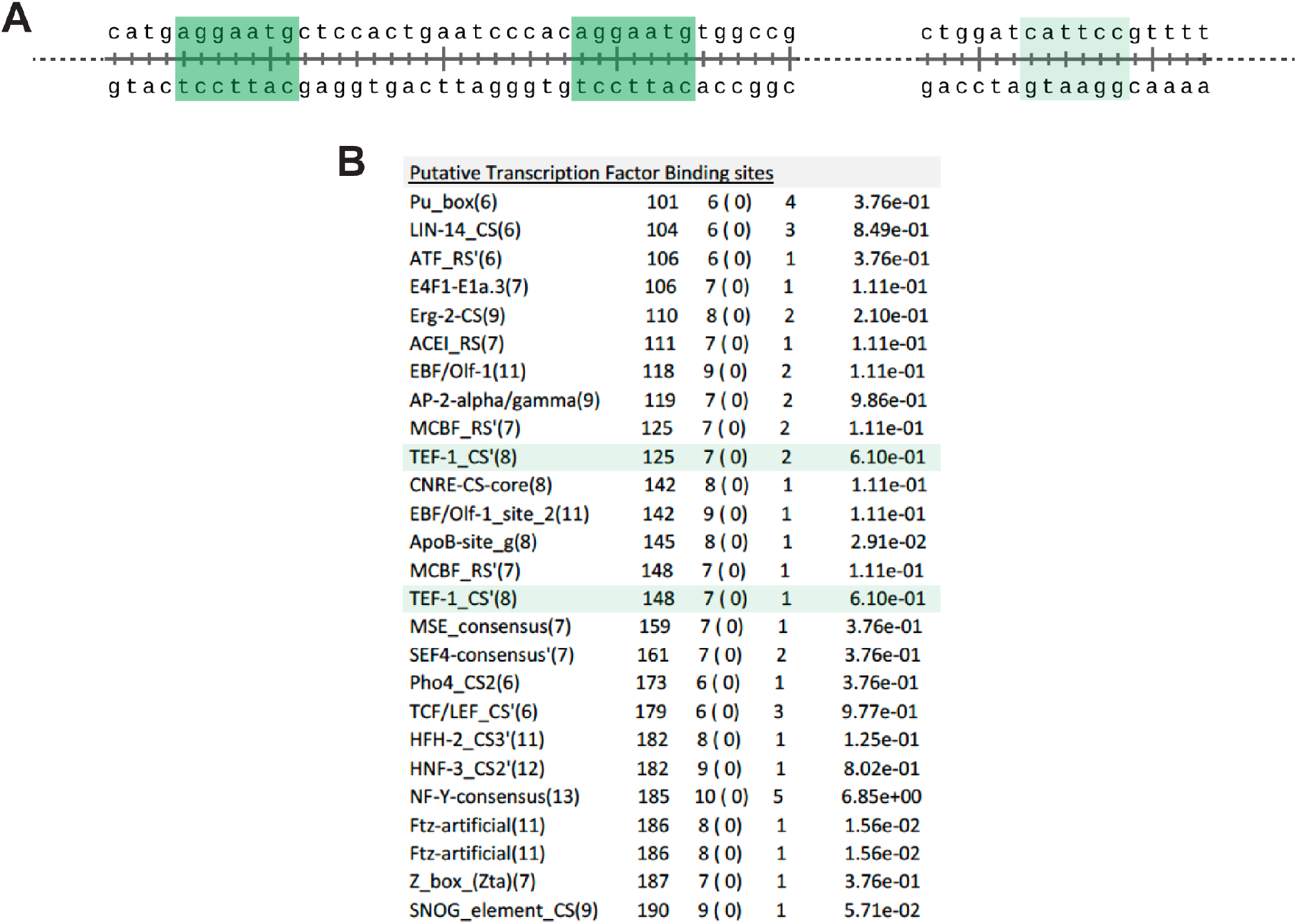
Transcription factor binding sites within the *1Kb-ctgfa* enhancer element. (A) A 190bp region was identified within the *1Kb-ctgfa* enhancer. Two perfect and one imperfect Tead binding sites are identified. (B) A full list of predicted transcription factor binding sites within the 190 bp *cis*-regulatory region.

**Figure S5.**
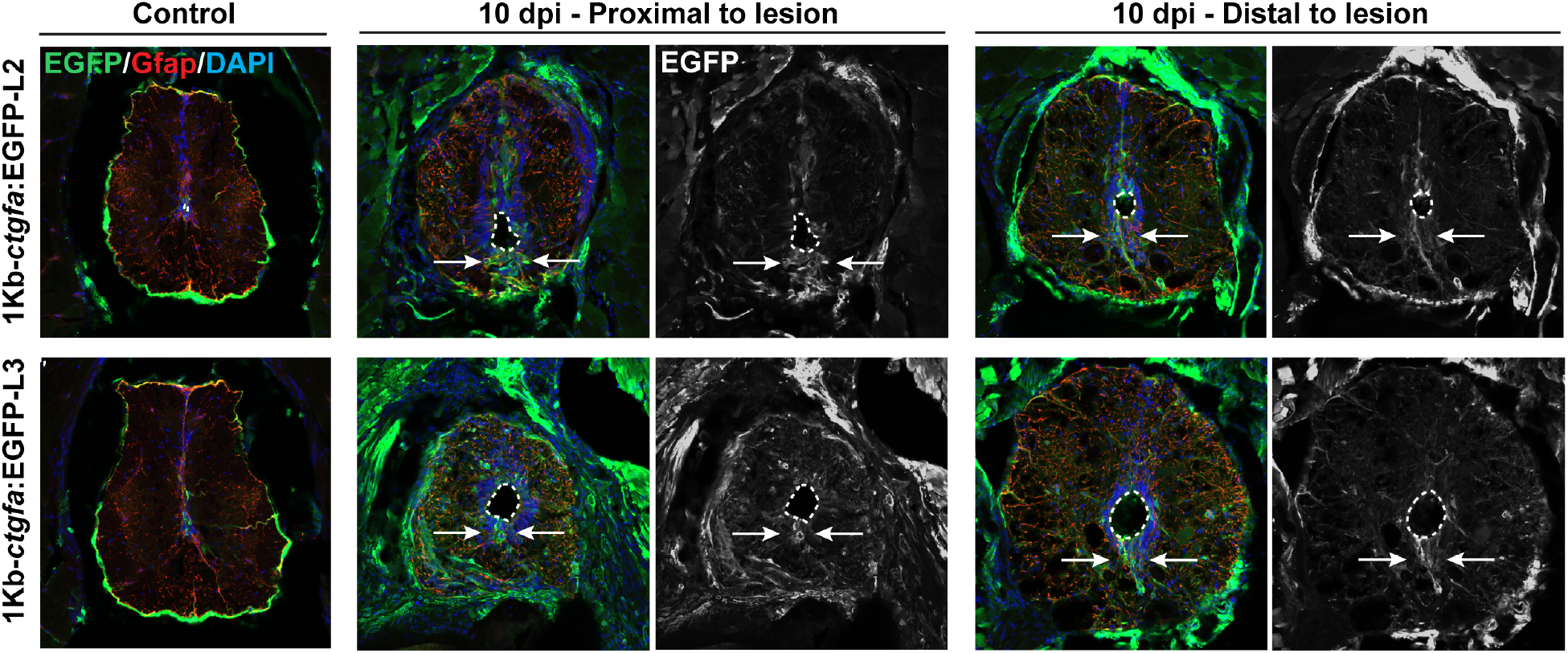
A 1Kb enhancer element directs *ctgfa* expression after SCI. EGFP and Gfap expression in two additional independent transgenic lines (L2 and L3) of *1Kb-ctgfa*:EGFP reporter at 10 dpi and in uninjured tissue. Shown are SC cross sections proximal (~100 μm) and distal (~500 μm) to the lesion core on the rostral side. Dotted lines delineate central canal edges. Arrows point to EGFP expression in ventral ependymal progenitor.

**Figure S6.**
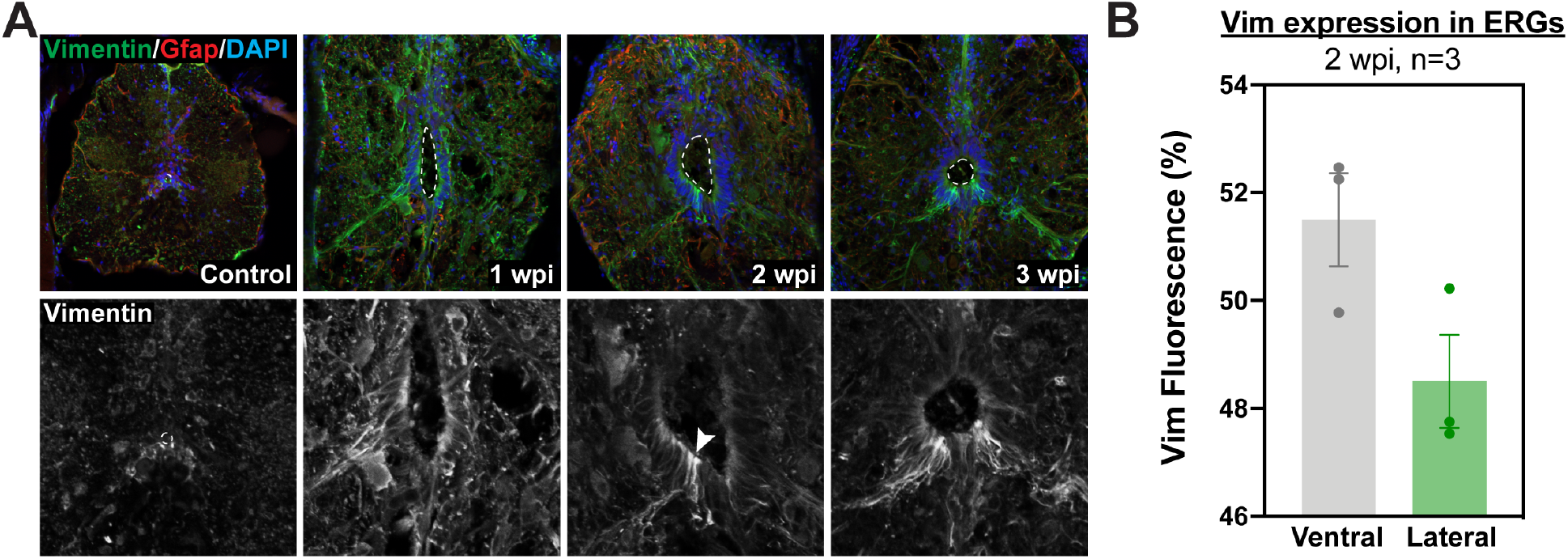
Vimentin expression after SCI. **(A)** Vim and Gfap immunostaining in wild-type zebrafish at 1, 2, and 3 wpi, and in uninjured controls. SC cross sections are shown. Dotted lines delineate central canal edges. Arrowheads point to Vim expression in ventral ependymal progenitors at 2 wpi. **(B)** For quantification, mean fluorescence was calculated in ventral and lateral ependymal domains at 2 wpi. Two sections per animals from 3 animals were used.

## Materials and Methods

### Zebrafish

Adult zebrafish of the Ekkwill strain were maintained as previously described. Male and female animals between 3 and 9 months of ~2 cm in length were used. Different manipulations (e.g. heat shock) can lead to different regeneration and recovery kinetics. Therefore, clutchmate control and experimental fish of similar size and equal sex distribution were used for all experiments. Spinal cord transection surgeries and regeneration analyses were performed in a blinded manner, and independent experiments were repeated using different clutches of animals. Newly constructed strains are described below.

Generation of transgenic *ctgfa*:mCherry zebrafish. The following primers were used to amplify a 5.5 kb genomic DNA region upstream of the *ctgfa* translational start site: ClaI forward primer 5’-atcgattttggctactttcagctaagactgg-3’ and ClaI reverse primer 5’-atcgattctttaaagtttgtagcaaaaagaaa-3’. The genomic fragment was cloned into PCR2.1-TOPO vector, then subcloned into ClaI digested PCS2-mCherry-Nitroreductase plasmid to generate *ctgfa*:mCherry-Nitroreductase clone. The clone was co-injected into one-cell stage wild-type embryos with I-SceI. Three founders were isolated and propagated. This line is referred to hereafter as *ctgfa*:mCherry, and the inducible Nitroreductase cassette was not used in this study. The full name of this line is Tg(*ctgfa*:mCherry-NTR). *ctgfa*:mCherry animals were analyzed as hemizygotes, or crossed into *gfap:*EGFP transgene (mi2002, [11]) to generate *ctgfa*:mCherry; *gfap:*EGFP animals.

### FACS sorting and RNA sequencing

Two mm SC sections, including the lesion site plus additional rostral and causal tissue proximal to the lesion, were collected from adult injured zebrafish at 5 dpi. Control tissue sections were collected from uninjured siblings. Wild type animals, *ctgfa*:mCherry, and *gfap*:EGFP animals were lesioned and dissociated as additional controls to set up FACS sorting gates. Tissues were dissociated using Trypsin and FACS sorted using a MoFlo Cell sorter. Total RNA was prepared using Tri reagent (Sigma). Total RNA was also directly prepared from two mm SC sections at 5, 10, 21 dpi, and from uninjured controls for bulk RNA-sequencing. TruSeq libraries were prepared in duplicate and sequenced on Illumina HiSeq 2000 using 50 bp single-end reading strategy. Quality QC and trimming of adapters and short sequences were performed using Fastx. Sequencing reads were then mapped to the zebrafish genome (Zv11) using Bowtie2, then assembled and quantified using the Cufflinks and Cuffdiff algorithms (22). RNA sequencing was performed and analyzed by the Bioinformatics Core at the Center for Regenerative Medicine at Washington University.

### Spinal cord transection and treatment

Zebrafish were anaesthetized using MS-222. Fine scissors were used to make a small incision that transects the spinal cord 4 mm caudal to the brainstem region. Complete transection was visually confirmed at the time of surgery. Injured animals were also assessed at 2 or 3 dpi to confirm loss of swim capacity post-surgery.

For recombinant CTGF treatment, sterile Gelfoam Absorbable Gelatin Sponge (Pfizer 09-0315-08) was cut into 1 mm3 pieces, soaked with either vehicle or recombinant CTGF peptides (HR-CTGF-CT, eBioscience, 14-8503-80), then cut into 4-6 smaller pieces. At 5 and 10 days postinjury, zebrafish were anaesthetized using MS-222. Fine scissors were used to make a longitudinal incision lateral and parallel to the spinal cord. Injured SC tissues were exposed without causing secondary injuries and gelfoam sponges were placed adjacent to the lesion site. The incision was closed and glued using Vetbond tissue adhesive material.

### Histology

Sixteen μm cross or 20 μm longitudinal cryosections of paraformaldehyde-fixed SC tissues were used. Tissue sections were imaged using a Zeiss AxioVision compound microscope (for *in situ* hybridization) or a Zeiss LSM 800 confocal microscope (for immunofluorescence).

*in situ* hybridization probes for *twist1a* were subcloned after amplification from 2 dpf zebrafish cDNA into PCR2.1-TOPO vectors (*twist1a* forward primer 5’-tgtgattgctctgctgttcc-3’, *twist1a* reverse primer 5’-ggtgaggcgattagcttctg-3’). Linearized vectors were used to generate the digoxygenin labeled cRNA probes. *in situ* hybridizations were performed as previously described (REF). Primary antibodies used in this study were rabbit anti-GFAP (Sigma, G9269, 1:1000), mouse anti-GFAP (ZIRC, Zrf1, 1:1000), and mouse anti-acetylated-α-tubulin (Sigma, T6793, 1:1000). Secondary antibodies (Invitrogen, 1:200) used in this study were Alexa Fluor 488, Alexa Fluor 594 goat anti-rabbit or anti-mouse antibodies. GFAP immunohistochemistry was performed on serial longitudinal sections. The diameter of the glial bridge and the diameter of the intact spinal cord rostral to the lesion were measured using ImageJ software. Bridging was calculated as a ratio of thesemeasurements. For each fish, percent bridging was calculated for the 3 longitudinal sections that show the most prominent bridging. Percent bridging was then averaged for each cohort. Mann Whitney tests were performed using Prism software to determine statistical significance between groups.

### Swim capacity assays

Zebrafish were exercised in groups of 8-12 in a 5L swim tunnel respirometer device (Loligo, cat#SW100605L, 120V/60Hz). After 5 minutes of acclimation inside the enclosed tunnel, water current velocity was increased every two minutes and fish swam against the current until they reached exhaustion. Exhausted animals were removed from the chamber without disturbing the remaining fish, while others continued to swim. Swim time and current velocity at exhaustion were recorded. Results were expressed as means ± SEM. An unpaired two-tailed Student’s t-test with Welch correction was performed using the Prism software to determine statistical significance of swim times between groups.

### Axon tracing

Anterograde axon tracing was performed on adult fish at 4 wpi. Fish were anaesthetized using MS-222 and fine scissors were used to transect the cord 4 mm rostral to the lesion site. Biocytin-soaked Gelfoam Gelatin Sponge was applied at the new injury site (Gelfoam, Pfizer, 09-0315-08; Biocytin, saturated solution, Sigma, B4261). Fish were euthanized 6 hours post-treatment and Biocytin was histologically detected using Alexa Fluor 594-conjugated Streptavidin (Molecular Probes, S-11227). Biocytin-labeled axons were quantified using the “threshold” and “particle analysis” tools in the ImageJ software (26). Four sections per fish at 0.5 (proximal) and 2 (distal) mm caudal to the lesion core, and 2 sections 1 mm rostral to the lesion, were analyzed. Axon growth was normalized to the efficiency of Biocytin labeling rostral to the lesion for each fish. The axon growth index was then normalized to the control group for each experiment. Mann-Whitney tests were performed using Prism software to determine statistical significance between groups.

